# Gut microbiome couples gut and brain during calorie restriction in treating obesity

**DOI:** 10.1101/2020.12.04.406579

**Authors:** Qiang Zeng, Qi Wang, Tianyuan Xiang, Lei Ou, Xiaoling Wu, Kaiye Cai, Chunyu Geng, Mo Han, Zhongxia Li, Zhonglin Li, Wen Wang, Tingting Yang, Fengyun Li, Huimin Ma, Xiaojuan Zhao, Na Mi, Hui Gao, Li Tong, Chi Zhang, Linyuan Wang, Bin Yan, Ziya Yu, Ziyu Wang, Canhui Lan, Xiaoning Wang, Yongli Li, Jun Wang

**Author notes:** Co First Authors: Qiang Zeng, MD, PhD., Qi Wang, PhD., Tianyuan Xiang, MD, PhD., Lei Ou, MD, PhD. Corresponding author: Jun Wang, PhD., Yongli Li, MD, PhD., Xiaoning Wang, PhD.

## Abstract

Calorie restriction (CR) has been widely recognized for its effect in reducing body weight and alleviating diabetes in humans, as well as prolonging life span in animal studies. Gut microbiome shifts contribute to part of the effects of CR, but little is known regarding their influences except on metabolism and immunity. Here we monitored gut microbiome using metagenomics and metatranscriptomics in obese individuals undergoing CR, and revealed microbial determinants that could contribute to successful weight loss. Microbiome changes are linked to changes in blood metabolome and hormones, which eventually correlate to brain functional changes as studied using functional magnetic resonance imaging (fMRI). Brain functional shifts indicate response of central neural system (CNS) to CR, and microbiome constitutes the keystone of gut-brain axis. Animal experiment further reaffirms the gut microbiome changes, metabolic and hormonal shifts of CR, while proteomic analysis of brain tissues suggest that epigenetic modifications of key proteins could explain responses of CNS to CR. Our study establishes linkage between CR, gut microbiome, metabolome/ hormones and CNS function, and demonstrates that CR has multi-facet, coordinated effects on the host, of which many could contribute to weight loss and other beneficial effects.

## Introduction

Calorie restriction (CR), i.e. designated reduction of food intake, has been shown to be an effective strategy in alleviating obesity and type II diabetes, yet current knowledge on the underlying mechanism is limited to a few studies mainly carried out in animal models^1–4^. Among them, gut microbiome changes were considered to play a central role, corresponding to the consensus that dysbiosis of gut microbiome is critical for the occurrence and development of obesity as well as obesity-related morbidities, including type II diabetes and hypertension^5, 6^. The shifts in microbiome under calorie restriction might synergize with the reduction of calorie intake by remodeling microbial metabolites and reduce energy contributions to host requirements, and at the same time increase beneficial metabolites including butyrate that alleviate obesity-caused inflammations and metabolic disorders^7^.

Another aspect of calorie restriction and related microbiome shifts, thus far largely ignored, is the behavioral changes in humans undergoing CR. Physiological changes under CR have been shown to contribute to reduction of ageing rate and oxidative stress, lowering risk factors for cardiovascular diseases and alleviate obesity, and changing endocrinal functions that correlate to metabolism and physical health^8^. Yet, CR is shown to be able to modulate behavioral changes in both human and experimental models, in measurements that include both objective (for instance psychological well-being, cognitive functioning) and subjective (mood, subjective feelings of hunger etc., determined usually by questionnaires) criteria^9^. However, currently available studies that have behavioral measurements are largely descriptive, leaving the exact mechanisms to be addressed that correlate other aspects of CR, including shifts in gut microbiome and metabolome, as well as brain functional responses^10^.

The function of central neural system (CNS) is tightly associated to gut microbiome, a connection that is increasingly appreciated as the gut-brain axis^11^. Gut microbiome is key producer of diverse neurotransmitters and neurotoxins under pathological condition or dysbiosis^12,13^. Such molecules are carried by the blood circulation system to reach CNS, and then modulate the function or inflammation that eventually affect behavior^14^. Recent studies revealed novel route of microbiome affecting CNS functions, including transmitting inflammation signals via the vagus nerve to CNS and then feed back to the gut^15^; or short-chain fatty acids including acetate that activate parasympathetic nervous system and eventually affect pancreatic beta-cell function and consequently energy metabolism^16^. It is extremely likely that gut microbiome shifts under CR might also be connected and contribute to CNS functionality of the host, and understanding the mechanisms as well as consequences of behavioral changes could further increase the effectiveness of CR.

In our current study, we have followed 35 extremely overweight individuals in their course of CR, and comprehensively analyzed their gut microbiome using metagenomics and metatranscriptomics, metabolome under CR and CNS functionality via functional magnetic resonance imaging(fMRI). We find that brain functions underwent certain changes that were connected to gut microbiome and metabolomic shifts. Parallel animal experiments reaffirm the findings in human that CR lead to critical gut microbiome and metabolome changes, and the CNS functionality could be due to epigenetic modifications (acetylation, phosphorylation etc) of key proteins. Our study reveals the multi-facet nature of CR on the host and deepens our understanding on the connectiveness of CNS function to CR and gut microbiome.

## Results

### CR leads to physiological improvements, metabolic shifts and brain function changes

In our study, 30 individuals with body mass index (BMI) > 28kg/m^2^ went through a designated scheme of CR. Within a step-wise reduction of calorie intake, the probands have their total calorie intake reduced to 2/3, 1/2, 1/3 and 1/4 of the normal diet every second day, while the other day they maintain their normal diet scheme, each step lasted 8 days. Fecal samples were at time point of before CR (baseline), then after 1/2 calorie diet (midpoint) and 1/4 calorie diet stage (end point) for metagenomic and metatrancriptomic analysis, accompanying serum were taken for metabolome, cytokine analysis as well as hormones that were related to metabolic health and obesity, and finally brain functions were analyzed with regard to regional homogeneity (Reho) using fMRI (**Figure S1**). After CR period ended, the probands were followed up for one month and fecal samples, serum and fMRI were measured after one month of relaxed control (post CR).

During CR, 25 out of 30 probands showed significant improvements in physiological conditions, defined as having >5% reduction in body weight and termed effective group (EG), the percentage is roughly consistent with previous findings regarding CR^17^. In the 25 EG individuals, physiological improvements include several physical characteristics and metabolic profile body, e.g. decreased in body mass (BM), body mass index (BMI), Waist Circumference(WC), Body Fat(BF), Systolic Blood Pressure(SBP), Diastolic Blood Pressure(DBP), Fasting Plasma Glucose(FPG), Glycosylated Hemoglobin (HbA1c), Total Cholesterol (TC), Glutamyl Transpeptidase (GGT) (**Table S1**); while those remained largely unchanged in the five individuals with unchanged body weight (ineffective group, IG).We found that compared with baseline, WC, FPG and HbA1c have significantly decreased at mid-piont (*p*< 0.05). Moreover, the WC and FPG continued to decrease till the time of end-point, while BMI, DBP and TG also declined compared with the baseline (*p* < 0.05). Furthermore, compared with baseline, BF, SBP, GGT have significantly decreased at post-CR (*p* < 0.05). There was no significant difference in sex, age, height, weight and metabolism parameters between the EG and IG.

Metabolome and hormone in the serum samples of EG group further revealed consequences of CR. Compared with baseline, the level of adiponectin (ADP), free fatty acid (FFA), VIP and caproic acid that have increased while leptin, propionic acid and isobutyric acid that have decreased at midpoint (*p* < 0.05). Further, adiponectin, FFA and dopamine continued to increase at end point, and leptin, somatostatin, acetic acid, propionic acid, isobutyric acid, butyric acid, isovaleric acid, valeric acid and caproic acid decreased significantly compared to baseline (*p* < 0.05). Eventually for post CR, TNF-a, adiponectin, FFA and acetic acid remained elevated while leptin, monocyte chemotactic protein 1(MCP-1) propionic acid, isobutyric acid, butyric acid, isovaleric acid, valeric acid and caproic acid remain significantly lower (*p* < 0.05) (**Table S2, Figure 1**).

**Figure 1:**
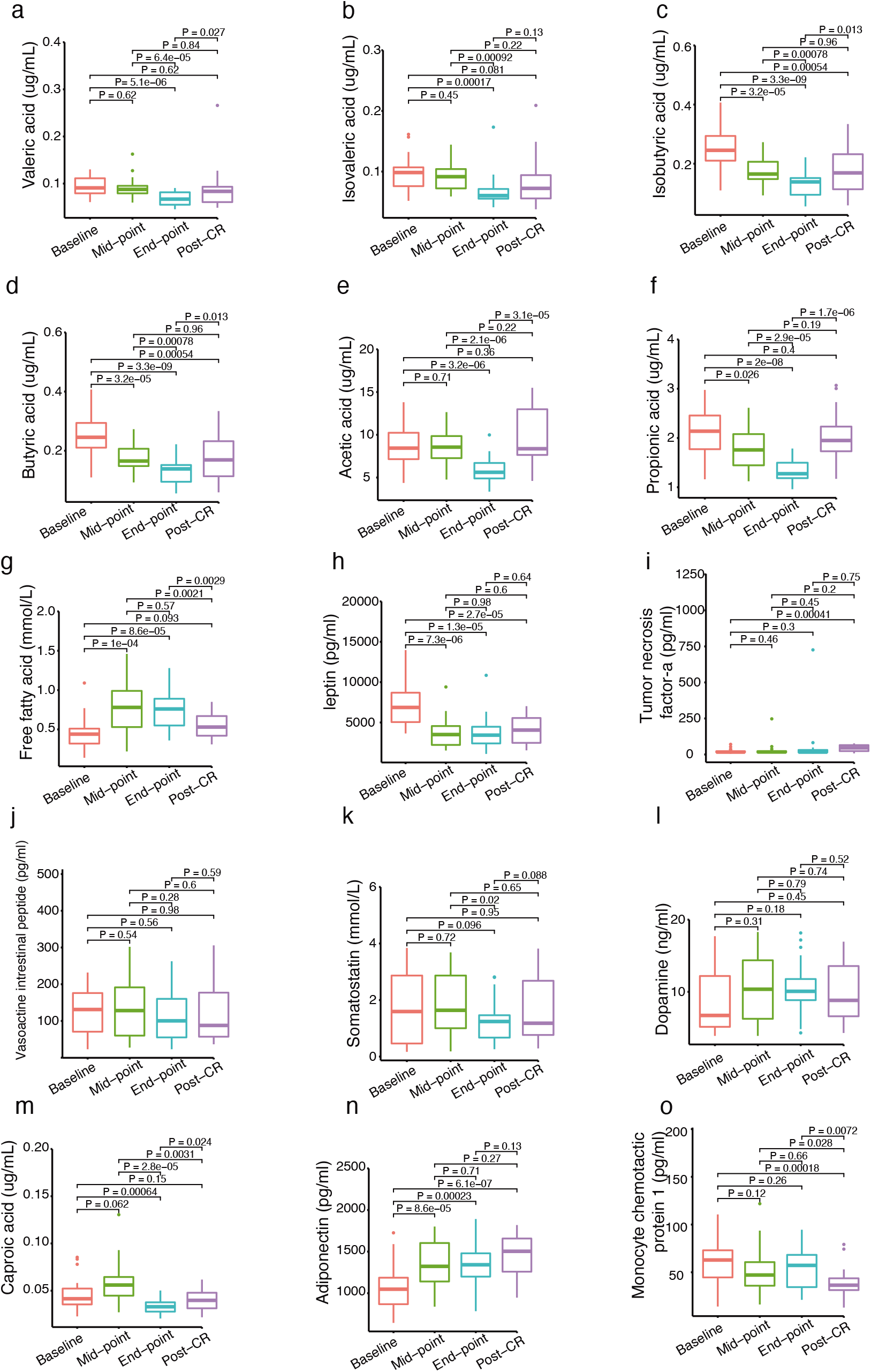
Summary of metabolite and hormone with significant changes during CR in effective group. a-o: different metabolites and hormones. Four time points were included, baseline: before CR; mid-point and end-point: middle CR and end of CR; post-CR: 1 month after the CR ended (Wilcoxon rank-sum test).

Accompanying the improvements in probands’ physiological health, brain functions of the participants undergoing CR showed significant changes as well. We compared brain function indicated by fMRI at baseline, midpoint, end point and post-CR covering ten areas of brain in the resting-state (**Figure S2**). Paired t test was used to analyze the obesity-related brain activity changes in resting-fMRI in time point-wise comparison in EG group. With GRF correction (q<0.005), we discovered that Reho of the left orbital inferior frontal gyrus decreased at midpoint compared with baseline, putamen decreased at end point compared with baseline, and right putamen, left dorsolateral prefrontal cortex, anterior cingulate cortex and right orbital inferior frontal cortex decreased at point post CR compared with baseline. We did not find statistically significant changes of brain activity in the reward circuits in pair-wise comparisons, nor in other brain areas (**Table S3, Figure S2**).

### Gut microbiome and metabolome shifts associated to brain functional changes

Gut microbiome in EG individuals underwent CR demonstrated significant compositional and functional changes, as revealed by metagenomic and metatranscriptomic analyses. DNA-based metagenomic analysis, which focuses on the standing community composition and functional pathway abundances, revealed that during CR, the alpha diversity indices of gut microbiome showed an increase in species composition though not significant, and then dropped significantly at the end of one-month washout afterwards (**Figure S3a**), similar trend was observed while using genus level composition (**Figure S3b**).

To explore signatures of the gut microbiome during CR, we identified multiple species and pathways that displayed significant abundance differences. During CR, species belonging to *Escherichia coli* and *Prevotella copri* and *Streptococcus salivarius* had significantly decreased while *Faecalibacterium prausnitzii, Roseburia hominis, Eubacterium hallii, Eggerthella lenta, Clostridium leptum, Akkermansia muciniphila* and *Bacteroides thetaiotaomicron* increased their abundances (Kruskal–Wallis test, all p < 0.05, Figure 2); and at functional pathway level, several amino acid degradation pathways (serine degradation, lysine degradation II and alanine degradation I), SCFAs synthesis and production pathways (Acetate synthesis IV, propionate synthesis II, propionate production II), glutamate metabolic pathways (Glutamate degradation I, Glutamate synthesis II, glutamine degradation II, GABA synthesis I and GABA synthesis II) and lactaldehyde degradation decreased their abundances in majority of individuals, and butyrate production I, GABA degradation, sulfate reduction, acetate to acetyl-CoA, lactate production increased in contrast (p < 0.05, Table S4). For those species and functional pathways with significant shifts during CR, species like *Bacteroides thetaiotaomicron*, functional pathways like butyrate production I, GABA degradation and sulfate reduction remained similar in abundances compared to during CR, demonstrating long-term effect of CR, while the rest regained similar abundances to that of before CR **(Figure 2)**. In contrast in IG group, besides *Akkermansia muciniphila*, those species and functional pathways only showed slight changes in shifts during and after CR, compared to the abundances before; Thus microbiome shifts could indeed be associated with the effectiveness in body weight reduction during CR.

**Figure 2:**
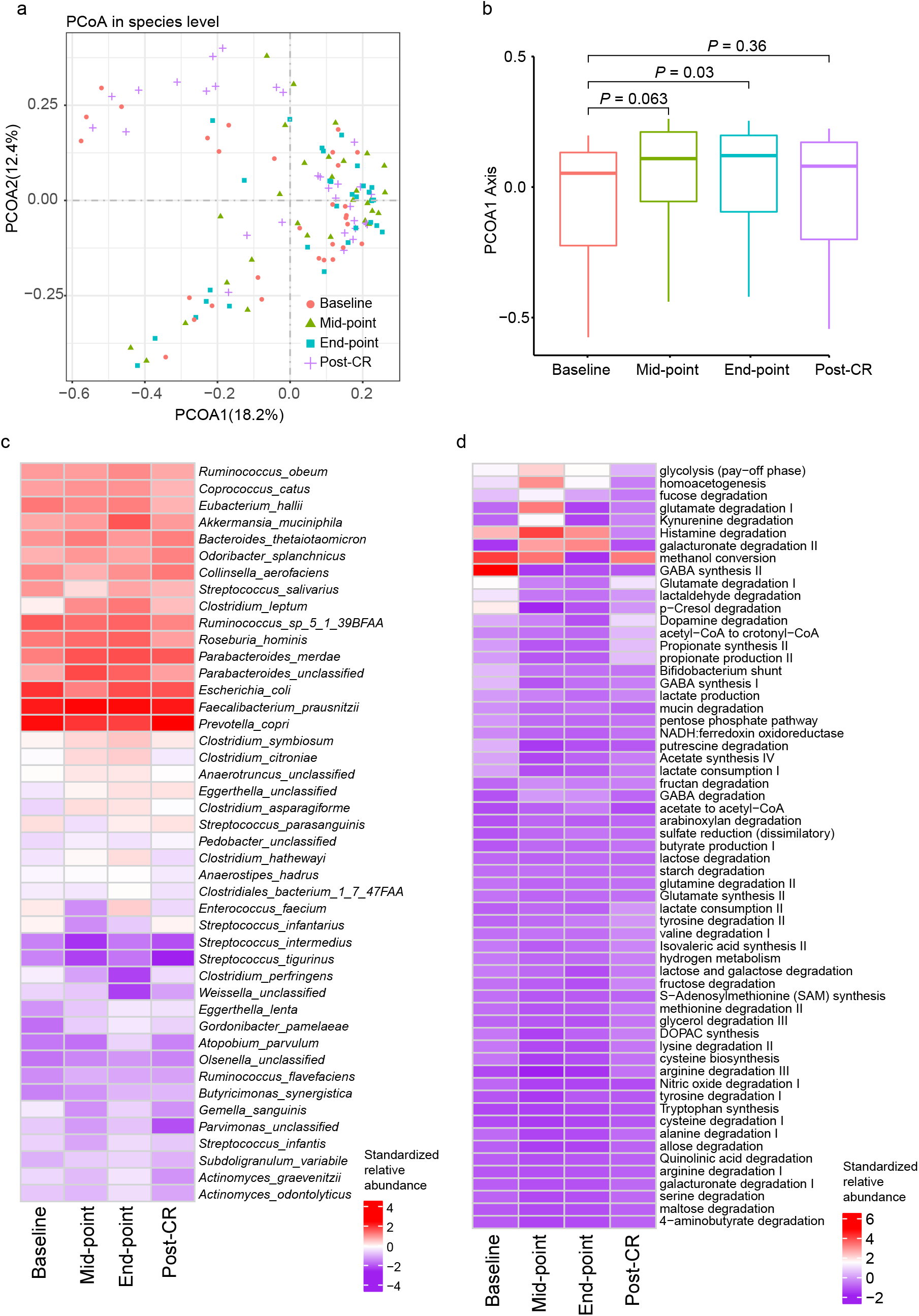
Summary of gut microbiome shifts CR treatment. A) The Principle Coordinate Analysis (PCoA) showing microbiome of all samples at different time points, PCoA was performed using Bray-Curtis dissimilarity at species-level. B) Coordinates of samples at different time points on PCOA1 axis, showing significant differences between baseline and post-CR (Wilcoxon rank-sum test, p< 0.05). Heatmaps present the average, standardized relative abundance of significantly changed (Kruskal-Wallis rank sum test, p < 0.05) species (C) and modules (D) versus four time points (see methods).

RNA-based metatranscriptomic analysis demonstrated functional pathways that are particularly active/inactive in the gut microbiome samples, and their dynamics during and after CR. For this purpose, we compared functional annotation results based on metagenomic and metatranscriptomic data. Overall, we found functional pathways like acetyl-CoA to crotonyl-CoA, allose degradation, serine degradation, alanine degradation I, propionate production II, lactate consumption I, maltose degradation, 4-aminobutyrate degradation, glutamate degradation, putrescine degradation, glutamate degradation III, asparagine degradation were under-represented in metatranscriptomic during CR, among those, acetyl-CoA to crotonyl-CoA, allose degradation, serine degradation, alanine degradation I, propionate production II, lactate consumption I, maltose degradation, 4-aminobutyrate degradation, glutamate degradation I, putrescine degradation were significantly changed in EG group, while in IG group those changes were not significant, demonstrating again the potentially important role of microbiome in shaping the outcome of CR **(Figure 2d)**.

Correlation analysis in EG group revealed a complex microbiome-metabolome/ hormone-brain function network **(Figure 3)**, and identified key microbial species and functional pathways which may contribute to the metabolites and hormones and finally lead to brain function shifts. We found a group of metabolites and hormones that might be influenced by the gut microbiota and were directly correlated to previously measured brain function changes. Putamen activitiy was negatively correlated with adiponectin, and was positively correlated with leptin and short chain fatty acids, such as propionic acid, isobutyric acid, acetic acid, and isovaleric acid. Notably, leptin was also positively correlated with right putamen, and isovaleric acid was positively correlated with right orbital inferior frontal cortex. Adiponectin was negatively correlated with right putamen and right orbital inferior frontal cortex. By extending the network, we eventually discover that the gut microbiota and their encoded pathways include amino acid synthesis, such as serine degradation, alanine degradation I and lysine degradation II are linked to brain function via the intermediate metabolites. For instance, putamen are connected in the network to *Faecalibacterium prausnitzii* by the latter’s positive correlation with lysine degradation II, and then positively correlated with leptin.

**Figure 3:**
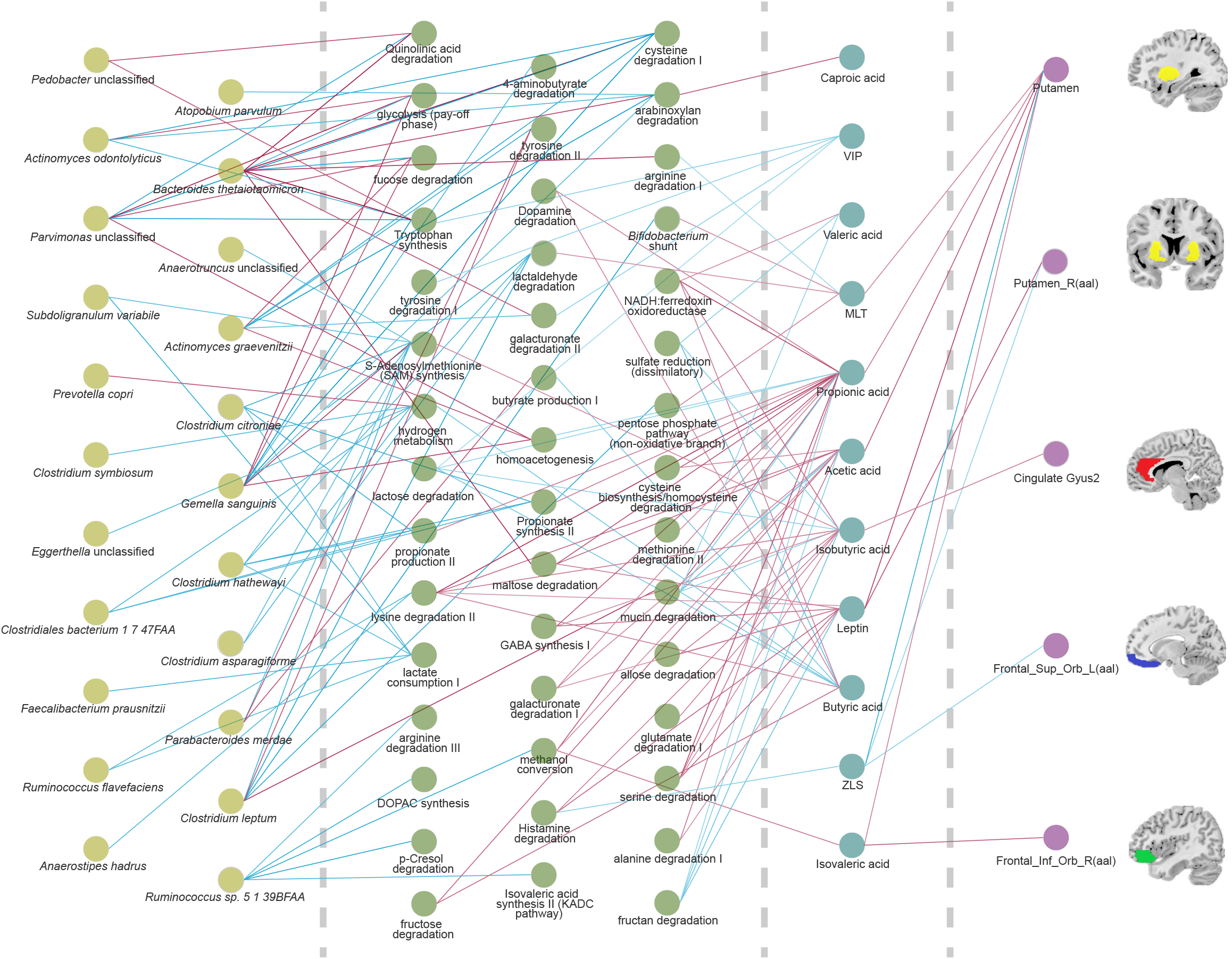
Correlational network of CR-changed bacterial species (first and second columns), functional pathways (third to fifth columns), metabolites and hormones (sixth column) and finally brain functional regions with significant changes under CR(Kruskal-Wallis rank sum test). Red lines denote positive and blue lines denote negative correlations (see results and methods).

### Animal experiment reaffirms microbiome and metabolome changes in CR

In order to reaffirm the correlations observed in human probands between CR and gut microbiome shifts, metabolism improvements as well as brain functional changes, especially investigating mechanisms underlying the brain functional changes, we replicated the CR in obese rat models. In high fat diet-induced obese rats, we randomly selected 17 and subjected rats to CR for a period of 16 days with similar scheme of human participants described above (2/3, 1/2, 1/3 and 1/4 daily dietary calories and each stage four days, but continuously in contrast to human study), another eight obese rats were used as control (obesity group). During the entire CR process, 14 rats had weight loss over 20% and were selected to replicate the EG group in humans, 2 failed (weight loss less than 20%) and 1 died **(Figure 4A)**.

**Figure 4.**
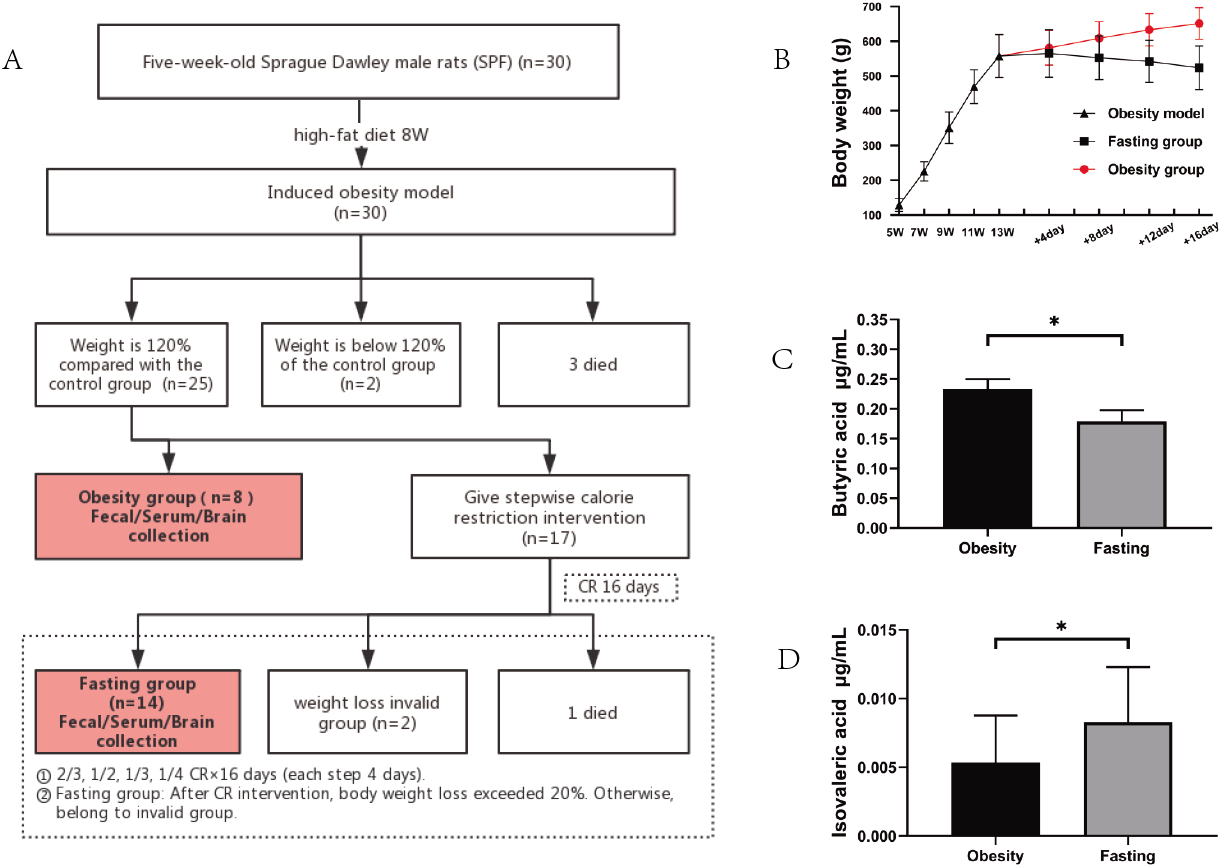
Experimental setup (A) and physiological shifts (B-D) in animal study. Obesity model was established using high-fat diet, and 14 rats were subsequently subjected to CR to mirror human study. CR lead to significant body weight loss, reduction in butyric acid and isovaleric acid in comparison to obesity rats (Wilcoxon rank-sum test) (see results and methods).

Metabolome analysis and hormonal measurements also reaffirms largely the changes observed in humans underwent CR. We discovered overall metabolome differences in comparison to that of CR rats, including that butyric acid decreased and Isovaleric acid increased in relative abundances (p<0.05). Moreover, adiponectin, leptin, dopamine and free fatty acid also showed increases during CR, in contrast to decreased TNF-α and MCP-1. Among those, adiponectin, MCP-1, free fatty acid and butyric acid, valeric acid and caproic acid mirror the trends observed in humans, demonstrating the consistent effects of CR on humans and rat models in modulating metabolome and hormones **(Figure 4)**.

### Epigenetic modifications of key proteins underlie brain functional shifts

In order to understand the mechanisms of CR and microbiome shifts correlate and thus potentially contribute to brain functional shifts, we have further conducted proteomics analysis in brain tissues of Fasting (n=14) and obesity (n=8) groups rats. Since proteins are ultimately the functional effectors of biological activity in body cells, in the section that follows, we will attempt to explore the potential role of epigenetic modification of key proteins in brain function transformation. We quantitatively profiled global proteomes, phospho-proteomes and acetyl-proteomes of 2 groups samples by using TMT labeling, HPLC fractionation and LC-MS/MS analysis.

A total of 5930 proteins were identified, of which 5116 contained quantitative information. If the threshold of differential expression was 1.2x and t-test p-value < 0.05 was significant, we found that 41 proteins were up-regulated and 32 proteins were down-regulated in Fasting vs Obesity comparison group (**Figure 5**). Quantification of proteins revealed little shifts in relative abundances of proteins in the brain tissues, but in the results of phospho-proteomes and acetyl-proteomes modification, we have a pleasantly surprised to find that a total of 7413 phosphorylation sites on 2608 proteins were identified. The 2608 identified phosphoprotein accounted for about 50.98% of the total identified brain proteins. High throughput analysis revealed a total of 1998 unique acetylation sites on 816 proteins, and these 816 identified acetyl-proteins account for only about 15.95% of the total proteins. We focus on the phosphorylation modification comparison between the Fasting group and the Obese group integrated analyses revealed that nearly two hundreds proteins and sites were identified as differentially expressed, 149 proteins were upregulated and 198 sites, 145 proteins were downregulated and 172 sites (FDR<1%) **(Table S4)**.

**Figure 5.**
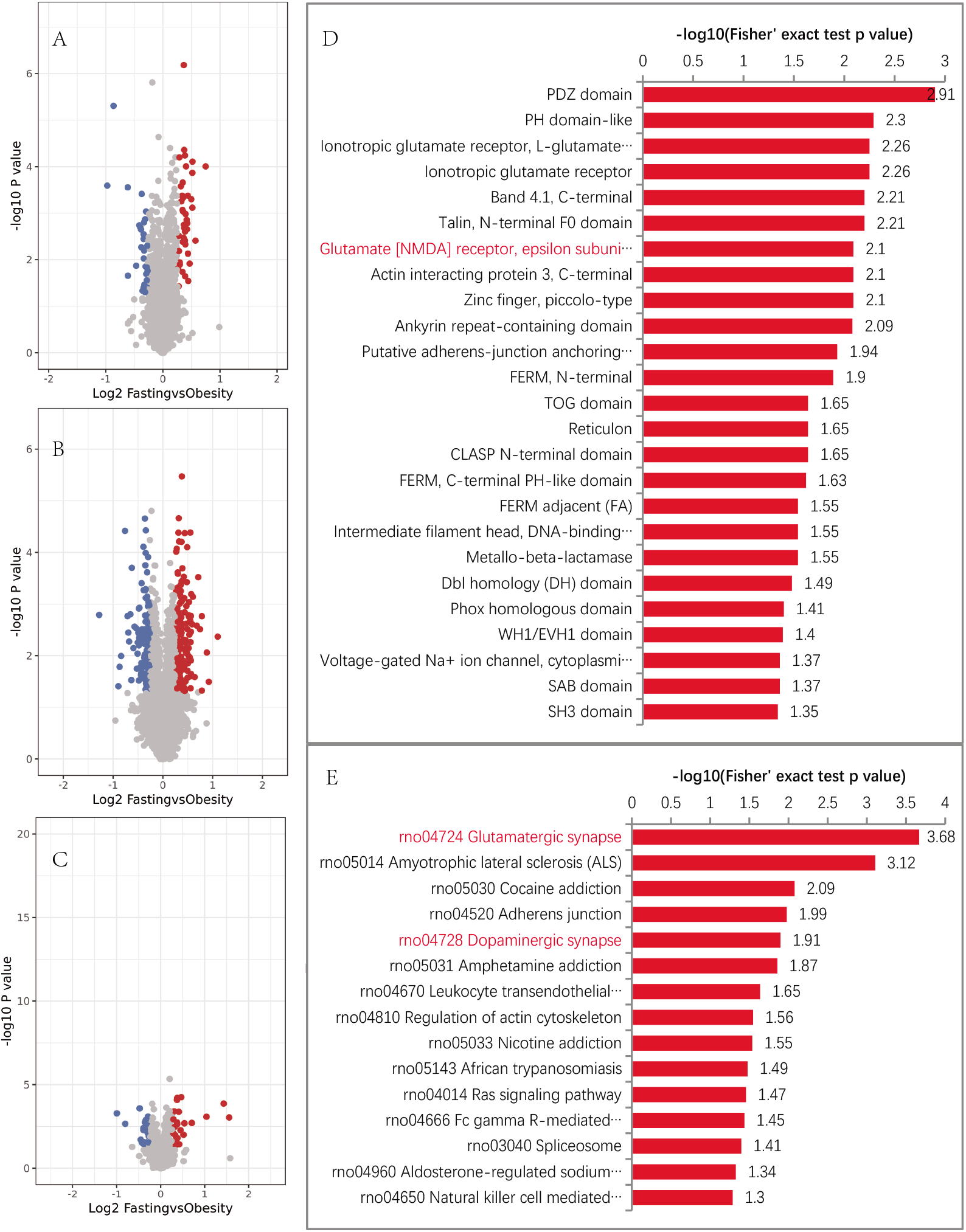
CR resulted in differences in rats’ brain proteome. Volcano plots show differences in global proteomes (A), phospho-proteomes(B) and acetyl-proteomes(C), each dot represents one protein, red dots show up-regulated proteins in CR rats while blue dots in obese rats. X-axis shows log2-transformed protein expression ratio change and y shows -log10 conversion of p-value (two-tailed Fisher’s exact test). Additional enrichment analyses show pathways with most differences between CR group and obese group, with (D) enriched based on KEGG pathway and (E) based protein domain (see results and methods).

The GO functional classification of the 2608 identified phosphorylated proteins was conducted based on their biological processes and molecular function. These phosphorylated proteins covered almost all cellular functions. With respect to biological processes, 188 and 90 respectively were involved in cellular and metabolic process proteins, 163 were involved in biological regulation (**Table S4**). Within the category of molecular function, the largest group (212 proteins) of GO annotated phosphorylated proteins was involved in binding activities, suggesting that phosphorylation is an important PTM in protein-protein interactions and in DNA transcription. The second largest group (59 proteins) has catalytic activity, indicating that the phosphorylation of enzymatic proteins may regulate or influence metabolic pathways (**Table S5**).

To further understand the differentially expressed proteins between fasting group and obesity group, functional classification and functional enrichment analyses (GO, KEGG pathway, and protein domain) were performed. Fisher’s exact test p-value (-log10[p-value]) was used to evaluate the enrichment levels of differentially expressed proteins; a larger p-value indicates more differentially expressed proteins enriched in this category. The enrichment analysis demonstrated that those proteins identified in the present study were mainly enriched in terms of functions in Glutamatergic synapse, Dopaminergic synapse, Regulation of actin cytoskeleton and Ras signaling pathway. Enrichment was also observed in proteins involved in Amphetamine addiction, Cocaine addiction and Nicotine addiction. All these results showed that the N-methyl-D-aspartate receptors (NMDARs) and glycogen synthase kinase3β (GSK3β) were indispensable and played critical role in those process (**Figure 5, FigureS4)**.

### Key correlations in our multi-omic study emphasize gut-brain axis

As extension of our findings from our multiple lines of observations, we then investigated the correlations we found between CR, gut microbiome composition and functional changes, metabolome and hormone and eventually brain function shifts, concluding that a large proportion supports the gut-brain axis. CR modulates several bacteria in human and/or rats that are known to be butyrate producers, including *Clostridium leptum, Eubacterium hallii, Roseburia hominis and Faecalibacterium prausnitzii*, or acetate producers such as *Clostridium hathewayi, Subdoligranulum variabile* and *Ruminococcus*; while another few were known to produce key neurotransmitter GABA, including *Streptococcus salivarius*. Accordingly, the metabolic pathways encoded by microbiome showed significant changes as well, as revealed by metagenomic and metatranscriptomic analyses, such as SCFAs synthesis and production pathways including acetate and butyrate production^18^, GABA synthesis. Despite relatively sparse studies on associations between gut microbiome and CNS function and majority of focus was their effect on metabolic and immunological diseases, we did find that some of the shifted bacteria were reported to be associated with CNS diseases as well, for instance *Akkermansia muciniphila* and *Prevotella copri* decreased in Parkinson’s disease in separate reports^19^, while *Clostridium symbiosum* and *Enterococcus faecium* have been found to be associated with schizophrenia^20^.

Short chain fatty acids and neurotransmitters affected by CR and partially produced by gut microbiome can directly enter circulation system and eventually enter CNS to affect brain functions, as established in previous studies. For instance, dopamine, Somatostatin and GABA are neurotransmitters that have critical roles in CNS functioning and implicated in various diseases including depression and anxiety, Parkinson’s disease and Alzheimers^21^. Short-chain fatty acids are known to modulate brain activities in recent years via multiple mechanisms, including modulate neuroinflammations by affecting immune cells (which in itself act notably via epigenic modulations on histone to affect certain gene expression), and stimulating other neurotransmitter/neuropeptides including 5-HT, GLP-1 and PYY in the intestinal tract^22^. In certain cases, metabolites do not only affect brain function, but can further drive brain functional feedbacks to other organs and change metabolic functions, such as acetate that changes brain activity under high fat diet and then transmit metabolic modulating signals to pancreas and digestive system via vagus nerve.

Eventually, proteomic analysis revealed proteins with epigenetic modification are involved in key process of brain and neural functions (Figure 6). One key group of proteins NMDAR are correlated to level changes in leptin, dopamine, TNF-a, and reported to influence the activity of and lead to brain nerve function changes; in particular, leptin has been shown to increase glutamatergic synaptogenesis in multiple brain regions, stimulation leads to increased phosphorylation of the NR2BY1472 residue and increases glutamatergic synaptogenesis, through enhancement of NMDARs function^23^; TNF-α is engaged in the abnormalities in NMDAR-mediated mEPSCs and learning erformance in anti-NMDAR encephalitis^24^, and finally adiponectin were protective against excitotoxicity induced by NMDARs in primary neurons^25^. In the CNS, NMDARs are known to participate in cell-cell communication, synaptic plasticity (memory, learning) and neuronal survival and death^26^,. By associating with various membrane receptors and extracellular or intracellular proteins in the complex, the NMDARs can contribute to various physiological processes such as learning and memory, and brain disorders such as stroke, schizophrenia and addiction^27^. Another protein affected by CR, GSK3β is a growth-signaling-sensitive kinase negatively regulated by inhibitory phosphorylation downstream of the insulin receptor, Wnt, and mTOR growth signaling pathways^28^. Notably, the role of GSK3β in phosphorylation of cytoskeletal proteins impacts neuronal plasticity, as cytoskeletal constituents are involved in the development and maintenance of neurites, and changes in the rate of stabilization/destabilization of microtubules could influence major cellular compartments of neurons, such as dendrites, spines, axons, and synapses. GSK3β has also been implicated in Alzheimer’s disease, where activation of GSK3β can promote tau hyperphosphorylation, neurofibrillary tangles, and amyloid plaques^29^.

## Discussion

Accumulating evidence from numerous studies asserted the critical roles of gut-brain axis in host health and diseases^30–32^. Commonly studied depression and anxiety are shown to associate with diseases including irritable bowel disease (IBS) and also obesity, although the causal relationships are still difficult to establish^33^. In both cases, gut microbial dybiosis was described to be one of the contributing factors, and even used as therapeutical target by approaches including probiotic treatment and fecal material transplantation^34, 35^. While in general population, the mental well-being is also established to be closely related to gut microbial contributions, in particular their potential of producing neuro-active substances^36^. CR has been used effectively in treating obesity in humans and many of the metabolic/immune aspects have been investigated^37^, yet our study elucidated the additional impact on central neural system, in particular the detailed regions of brains responsible for various perception, feedback and ultimately behaviors. Out of ten regions we inspected with fMRI, five showed significant changes due to CR at one time point, showing that CR could actually cast effect on the CNS activities and functions. More importantly we could conclude that certain changes in regions responsible for attention, learning, self-control and decision making (left dorsolateral prefrontal cortex, anterior cingulate cortex and right orbital inferior frontal cortex) are long-term, that the changes in them are significant one month after CR ended. It is highly likely that they would be able to lead to beneficial effect in terms of controlling food/calorie intake, and eventually benefit the obese patients that underwent CR by casting long-term effect, it has to be however carried out in future studies with more behavior analyses.

While microbiome shifts under dietary change, including CR are investigated before, we paid more attention to the consequent modulation of metabolome, cytokines and peptide hormones that are underlying the metabolic benefits of CR and eventually also contribute to CNS functional changes. Metagenomic analyses in human and rats indicate an overall shift in microbial community and functional potentials, accompanied by confirmation of significant changes in a collection of fatty acids, cytokines and hormones. In particular, C2-C6 fatty acids showed a general trend of decrease due to CR, which we conclude is mainly due to the general decrease of dietary fiber in combination with microbiome shifts and indicate less energy intake of the probands from food. Higher adipodectin and decreased leptin are indications of improved metabolism and weight loss in humans, but only adipodectin can be replicated in animal models, suggesting that it might be more essential to the effect of CR. Pro-inflammatory and pluripotent cytokine MCP-1 (also known as CCL2) was lowered by CR as well, but we again observed opposite trend for TNF-a between human and animal model. We established correlation between microbiome shifts and parameters mentioned above in our dataset, and while many of the associations can be supported by previous studies, their mechanistic explanations still require designated studies.

Using animal model, we investigated the mechanistic aspects of brain functional changes during CR, in particular we show that protein expression levels were not significantly altered during fasting, however the epigenetic modifications were. Proteomic analysis revealed overall changes in phosphorylation and acetylation, and two key cellular pathways were found to be with most changes during fasting: NMDARs and GSK3β. The NMDARs are associated with various membrane receptors^38^ and extracellular or intracellular proteins and contribute to various physiological processes^39^ such as learning^40^ and memory^41^, and brain disorders such as stroke^42^, schizophrenia^43^ and addiction^44^; while GSK3β affects phosphorylation of cytoskeletal proteins and impacts neuronal plasticity^45^, and eventually influences stabilization/ destabilization of microtubules in major cellular compartments of neurons. Such epigenetic changes have been reported to be associated with specific metabolites, cytokines and hormones, which in our study originate or associated with gut microbiome shifts. And we could translate such associations to the findings in humans, that specific brain activities were modulated by such microbiome-metabolites-protein axis. Moreover, many of the changes in brain functional activities maintained after CR ended, and could point to longer-term behavioral changes that will prolong the metabolic benefits of the probands.

In conclusion, we carried out CR in obese individuals and lead to weight loss and improvements in metabolism, and associated microbiome shifts contributed to metabolome, cytokine and hormonal changes, which eventually correlated with brain functional changes. We also investigated the molecular mechanism of gut-brain connections and determined primarily that the key contribution came from protein epigenetic changes. Our study deepens the understanding into CR and connects the desired brain activity and behavioral changes via gut-brain axis, an aspect that entices future investigations.

## EXPERIMENTAL MODEL AND SUBJECT DETAILS

### Ethical compliance

Ethical approval was obtained from the Ethics Committee at the Henan Provincial People’s Hospital, People’s Hospital of Zhengzhou University and the Ethical Committees of the Zhengzhou People’s Hospital. The study protocol was registered at https://clinicaltrials.gov (Protocol ID: 20180520). The study design complied with all relevant ethical regulations, aligning with the Helsinki Declaration and in accordance with privacy legislation. All participants provided written informed consent.

## EXPERIMENTAL DESIGN

### Human Subjects

The study was conducted at the Henan Provincial People’s Hospital, People’s Hospital of Zhengzhou University. We initially recruited 41 right-handed volunteers with obesity, and the inclusion criteria: participants in the recruited BMI range (28 to 45 kg/m^2^) who had hypertension, diabetes, hyperlipidemia, hypeluricemia, metabolic syndrome, but these diseases were well-controlled. Exclusion criteria were serious cardiopathy, liver dysfunction, renal disease, hypoglycemia, anemia, malnutrition, hemopoietic system disease, systemic immune system disease, infectious disease, and neurological disorders. One candidate was excluded from the study due to polycystic ovarian syndrome ultrasound examination. Subject who cloud not undergo MRI due to claustrophobia, implanted metal, pregnancy was excluded (n=1). In addition, four volunteers were excluded because of too much body fat that prevented them from fitting into the magnetic resonance coil comfortably, and abnormal anatomical structure in ectocinerea. Therefore, 35 subjects with obesity (BMI 35.29±4.15kg/m^2^; age36.26±9.14 years; 16 female). Two nutritionists made the individualized ketogenic calorie-restricted diet via the dietary records of normal diet of every volunteers. The prescribed diets carried out according to 2/3, 1/2, 1/3 and 1/4 of the original normal caloric intake, with 36% carbohydrate, 52% fat, 12% protein. Multivitamins and mucopolysaccharides were recommended, as was 30 min of walking or swimming per day. The study was approved by the ethic committee of Henan Provincial People’s Hospital. All participants were fully informed of the study procedures and signed informed consent to participant in the study.

35 participants with obesity were randomly divided into seven groups of five. Restricted diet experiment procedures consisted of three stages. The first stage was normal diet with no restriction on calories and food types, which lasted for four days.At this stage, subjects should fill in dietary records, which included the type and weight of their diet. The second stage was restricted diet of high control, and we adopted restricted diet every other day method.At this stage fasted diet was carried out according to 2/3, 1/2, 1/3 and 1/4 of the original normal caloric intake. Each caloric stage last for 7-8 days, and the total cycle of fast diet was 31 days. The third stage was restricted diet of low control and started after the intervention of the second stage for a month.Subjects were allowed to continue the diet fasted diet every other day. It was suggested that the daily caloric intake was 600kal/day for males and 500kal/day for females. Participants underwent data collection on four occasions;first prior to initiation of the diet program, after 1/2 calorie diet and1/4 calorie diet stage, then after one month of low control.The data collection consisted of physiological and biochemical index, some scales, feces, blood, and MRI data.Subjects completed the Restricted Diet Scale (RS), the Dutch eating behavior questionnaire (DEBQ) and Three-factor Diet Questionnaire (TFEQ) to assess differences in eating behavior.Sleep quality was assessed using Pittsburgh Sleep Quality Index (PSQI), anxiety and depression levels using Hamilton anxiety scale (HAMA) and Hamilton depression scale (HAMD).The physiological and biochemical index contained the analysis of human body composition, blood glucose, blood pressure, blood lipid, hepatorenal and kidney function, C-Reactive Protein and food intolerance test. The detailed information of MRI is introduced in the following.

### Animals

All experiments were approved by the Animal Care and Use Committee of the Zhengzhou People’s Hospital and were performed in accordance with institutional guidelines. Five-week-old Sprague Dawley male rats (SPF) (n=30) were used in this experiment, one per cage, in an environmental-controlled room (23-24 °C, 60% humidity). At the beginning of the experiment, rats were allowed ad libitum access to water and synthetic diets. The model rats were given high-fat diet (HFD) containing 35.8% fat, 20.7% protein, and 35% carbohydrates (n=30, D12492; Research Diets, ChangZhou, JiangSu, CHINA). Body weight and diet consumption were recorded once every two days. After 8 weeks of feeding, 25 rats were identified as obese rats while the body weight of the high energy diet group was 20% higher than average body weight in the maintenance diet control group (402 ± 17.56 g, control vs. 557.64 ± 12.35 g, obesity; *P*<0.05). For high-fat-diet experiments, 5 rats were excluded from the study: 2 rats failed to establish obesity model and 3 were died.

In order to reaffirm the correlations observed in human probands between CR and gut microbiome shifts, metabolism improvements as well as brain functional changes, we replicated the CR in obese rat models. In total 25 obese rats, 17 were randomly selected and in total underwent CR for a period of 16 days like participants described above (2/3, 1/2, 1/3 and 1/4 daily dietary calories, each phase four days), 14 rats had increased weight loss over 20% and was selected to replicate the EG group in humans (called Fasting group), 2 failed, and 1 died. Another 8 obese rats were used as control (called Obesity group). During this time, body weight, the brain-gut peptide (MCP-1, TNF-a, Adiponectin, Free fatty acids, Dopamine, Leptin) and SCFAs (Short-chain fatty acids, including acetic acid, propionic acid, butyric acid, valeric acid, isobutyric acid, isovaleric acid, caproic acid) in serum were measured and compared between the three groups. Rats were then euthanized, brains were taken and subjected to proteomics analysis. Serum, feces, and organs were collected for downstream analysis. Animals were humanely killed, and all the experiments were conducted in full compliance with the strict guidelines of the Zhengzhou People’s Hospital policy on animal care and use.

## METHOD DETAILS

### Hormone Measurements

Participants’ blood samples were collected on the morning after fasting 8 hours. The first time was on the baseline, the second time was at the end of 1/2 calorie diet of the original normal caloric intake, the third time was at the end of 1/4 calorie diet of the original normal caloric intake, the fourth time was at the end of restricted diet of low control. All samples from each participant weremeasured with in the same assay and were centrifuged as soon as possible to obtain serum, which were then stored in a minus 80 degree refrigerator until analysis.All the brain-gut peptide in serum were measured by ELISA (MULTISKAN MK3, *Thermo, USA*) which including Leptin, MCP-1, TNF-a, IL-6, IL8, Adiponectin, Free fatty acid,

LPS, LPL, Cb-1, 5-HT, VIP, Somatostatin, Melatonin, IGF-1, DA, PPAR-γ and BDNF.

### fMRI Data Acquisition

The MRI data includes anatomical MRI, resting-state fMRI. T2-weighted dark-fluid and T2 weighted MR images were acquired to exclude brain structural abnormality. The resting-state fMRI data were collected to observe the effects of CR on the brain during rest. The subjects were instructed to keep their eyes closed, not to fall asleep and keep their brain empty during the resting-state fMRI data acquisition. Resting-state data were acquired using a prototype simultaneous multi-slice echo planar imaging (SMS-EPI) sequence with the following parameters: TR = 2000 ms, TE = 35 ms, FOV = 220 mm × 220 mm; matrix size =94 × 94, slices = 75, slice thickness = 2.2 mm, flip angle = 80°, and SMS factor = 3.

### fMRI data Pre-processing

The resting-state fMRI data were analyzed using DPABI (http://rfmri.org/dpabi) and SPM12 (www.fil.ion.ucl.ac.uk/spm). Data preprocessing included the following steps. At first, DICOM format data were converted to NIFTI format. Second, the 10 initial scans of all experiment runs were discarded as a result of magnetic equilibration effects and the scanning noise adapted by participants. Third, the data were slice-timing and head movement corrected. Fourth, all functional datasets were normalized to the Montreal Neurological Institute (MNI) standard space by linearly registering, using echo-planar imaging (EPI) template. Fifth, all datasets were smoothed using a 6 mm full width at half maximum (FWHM). Sixth, the data were detrended to eliminate the linear trend of time courses and filtered with low frequency fluctuations (0.01–0.08Hz). Finally, six head motion parameters, white matter mask, the whole-brain mask and cerebrospinal fluid mask were regressed out of the EPI time series^46^.The results of pretreatment were as follows: five subjects were excluded due to excessive their six head movement parameters (displacement > 2 mm and/or rotation > 2 degrees). The final 30 cases met the standard.

### Metagenomic and metatranscriptomic sequencing

DNA was extracted from fecal samples using Magen Magpure Stool DNA KF Kit B (Magen, China) according to the manufacturer’s instructions. Construction of a paired-end library with insert size of 250bp was performed according to the manufacturer’s instruction, and the DNA library was sequenced with PE reads of 2*100bp on the BGISEQ-500 platform (BGISEQ, CN).

RNA was extracted from fecal samples using Qiagen RNeasy PowerMicrobiome Kit (Qiagen, Germany) according to the manufacturer’s instructions. Construction of a paired-end library using MGIEasy RNA Library Prep Set (MGI, China) with insert size of 250bp was performed according to the manufacturer’s instruction, and the RNA library was sequenced with PE reads of 2*100bp on the BGISEQ-500 platform (BGISEQ, CN).

### Metagenomic and metatranscriptomic sequencing

DNA was extracted from fecal samples using Magen Magpure Stool DNA KF Kit B (Magen, China) according to the manufacturer’s instructions. Construction of a paired-end library with insert size of 250bp was performed according to the manufacturer’s instruction, and the DNA library was sequenced with PE reads of 2*100bp on the BGISEQ-500 platform (BGISEQ, CN).

RNA was extracted from fecal samples using Qiagen RNeasy PowerMicrobiome Kit (Qiagen, Germany) according to the manufacturer’s instructions. Construction of a paired-end library using MGIEasy RNA Library Prep Set (MGI, China) with insert size of 250bp was performed according to the manufacturer’s instruction, and the RNA library was sequenced with PE reads of 2*100bp on the BGISEQ-500 platform (BGISEQ, CN).

### Quality control and host genome filtering of Metagenomic and metatranscriptomic reads

The raw reads that had 50% low quality bases (quality ≤ 20) or more than five ambiguous bases were excluded. The remaining reads were mapped to the human genome (hg19) by SOAP v2.22^47^ (-m 100 -x 600 -v 7 -p 6 -l 30 -r 1 -M 4 -c 0.95,), and the matching reads were removed^48^. The high-quality non-human reads were defined as cleaned reads.

### Acquisition of gene abundance and taxonomic profiles from metagenomic samples

The cleaned reads were aligned against the latest 11.4 M human gut microbial gene catalog^49^ through SOAP v2.22 (-m 100 -x 600 -v 7 -p 6 -l 30 -r 1 -M 4 -c 0.9) to generate the gene abundance profile. To obtain the taxonomic profiles, metaphlan2^50^(--input_type fastq --ignore_viruses --nproc 6) was used to generate phyla, genera, and species profiles from the clean reads.

### Calculation of gut microbiome functional profiles

Putative amino acid sequences were translated from the gene catalog^49^ and aligned against the proteins or domains in the KEGG databases^51^ (release 79.0, with animal and plant genes removed) using BLASTP (v2.26, default parameter, except -m 8 -e 1e-5 -F -a 6 -b 50). Each protein was assigned to a KEGG ortholog group on the basis of the highest scoring annotated hit(s) containing at least one segment pair scoring over 60 bits.

The relative abundance profile of KOs was determined by summing the relative abundance of genes from each KO using the mapped reads per sample^49^. The abundance of each gut metabolic module (GMM) (-a 2 -d GMM.v1.07.txt -s average) and gut neuroactive module (GBM) (default parameter) were calculated as shown in the former article^52, 53^.

The gut microbiome functional profiles were calculated by metatranscriptomic functional profile divide Metagenomic functional profile.

### Proteomic Profiling

The global proteome and phospho-proteome were processed according to adapted protocols from our previous studies (Huang et al., 2017; Mertins et al., 2016). In brief, cryo-pulverized brain tissue from each rat was lysed at 4°C using 8M urea lysis buffer. Extracted proteins were reduced using dithiothreitol and alkylated with iodoacetamide before digestion using LysC for two hours followed with trypsin overnight. Both digestion steps were performed at a 1:50 enzyme:protein ratio. For relative quantification of the global proteome and phospho-proteome by liquid chromatography tandem mass spectrometry (LC-MS/MS), 400 mg per patient, as measured on protein level (BCA protein concentration determination kit; before digestion) was labeled with 10-plexing tandem mass tags (TMT-10; Thermo Scientific) following the manufactures instructions. In parallel to the global proteome, phospho-proteome, and acetylome, an additional 500ug of TMT-labeled peptides per patient were enriched for phosphotyrosine peptides using phospho-tyrosine antibodies and analyzed as a single fraction on the mass spectrometer. All proteomic based data were collected on a Lumos mass spectrometer (Thermo Fisher Scientific) and the resultingspectra weresearched usingSpectrum Mill (Agilent, version12.212).

## QUANTIFICATION AND STATISTICAL ANALYSIS

### Hormone Analysis

We counted the content of brain-gut peptide in the serum of participants at four different time points. Firstly, we checked whether the four groups of data conform to the normal distribution or not. I fall the four groups of samples accorded with the normal distribution which could be described by mean number and the variance equality, they could be tested by single factor variance analysis (ANOVA) and a 2-tailed test with P≤0.05 would be regarded as statistically significant. If the four sets of data did not conform to normal distribution, we would choose non-parametric test, such as Mann-Whitney, Kolmogorov-Smirnov, and Wald-Wolfowitz test. The analyses were performed using SPSS version 17.0 (SPSS Institute, Inc).

### Analysis of fMRI data

Calculation of ReHo. Kendall’s coefficient concordance (KCC) can measure the similarity of a number of time series. A method was developed based on the regional homogeneity (ReHo) and described by Zang et al. (2004) The ReHo maps were used to quantitatively analyze the consistency of each voxel in brain area and the neighboring voxel (6, 18, and 26, respectively) activity level. It can describe the local characteristics of brain spontaneous activity^54^. All the four sessions’ ReHo maps were collected from the resting-state. The paired t-test was calculated to compare different conditions between two sessions’ ReHo values (GRF, voxel P<0.005 corrected), saved the GRF results as a mask, and generated a brain area activation report to see if there were brain areas in the peak in the four networks, including reward circuit, learning and memory circuit, inhibitory control circuit and sense and drive circuit^55^. The selection principles of the (Region of Interest, ROI) in these networks are as follows: Firstly, we set the peak point of the paired t-test results which were included in the four networks as the center of ROIs. Secondly, the brain regions that were not included in the results of the paired t-test, but were important for obesity research, were selected. The selected brain regions in the four networks were stored as the mask of seed points by AAL (61×73×61) template, which was loaded into the results of the paired t-test, the activation of corresponding brain regions was obtained, and the peak coordinates and t values of the brain regions were found.

### Richness and diversity analysis

Alpha diversity (within samples) at species, genus and KO levels was quantified by the Shannon index based on the relative abundance profile. Beta diversity was calculated based on Bray-Curtis distance (R 3.2.5, vegan package 2.4-4).

### Differential analysis of the gut microbiome and metatranscriptomic

MWAS^56^ was used to investigate the differences in composition versus four stages. (Paired wilcoxon rank-sum test and Kruskal-wallis rank sum test, *P*<0.05 as cutoff.)

### The correlation of gut microbiome and metatranscriptomic to clinical index

The correlation coefficient was analyzed by Spearman’s rank correlation (p < 0.05). Then the software, Cytoscape^57^ 3.4.0, was used to visualize the network (abs(Spearman’s correlation coefficient)>=0.2, p < 0.05).

### Normalization of Proteomics Data

Relative expression data derived from proteomics profiling (proteome, phospho-proteome, acetylome) were separately normalized by sample using robust z-scores(z_r_): z_r_=(x-M)/MAD. In this expression, M is the median expression of the sample; and MAD is the median absolute deviation of the sample. To allow for a better comparison across data types, WNT samples were excluded from these analyses as they lacked all but the proteomic data, and there was insufficient high-quality tissue to perform the additional assays.

