## supplemental materials for "Gut microbiome couples gut and brain during calorie restriction in treating obesity"

**Table S1 Summary of clinical characteristics of our cohort in the study. Values are presented as Mean±SD. BW: Body weight; BMI: Body Mass Index; WC: Waist circumference; SM: Skeletal Muscle; BF: Body fat; PBF: Percentage of Body Fat; SBP: Systolic Blood Pressure; DBP: diastolic blood pressure; FPG: Fasting Plasma Glucose; HbA1c: glycosylated hemoglobin; TG: Triglyceride; TC: Total Cholesterol; LDL-C: Low Density Lipoprotein-Cholesterol; ALT: Alanine Transaminase; GGT: Glutamyl Transpeptidase; ALP: Alkaline Phosphatase.**

| roup / Demographic data | Total Population(n=30) | EG(n=25) | IG(n=5) | *p* |
|---|---|---|---|---|
| **Females n (%)** | 14 | 12 | 2 | 0.157 |
| **Age（years）** | 37.6±9.05 | 37.64±9.37 | 37.40±8.20 | 0.406 |
| **Height (cm)** | 166.87±8. | 167.12±8.29 | 165.6±9.48 | 0.554 |
| **BW(kg)** | 96.81±14.99 | 97.53±15.67 | 93.18±11.66 | 0.206 |
| **BMI(kg/m$^2$)** | 34.54±3.84 | 34.64±4.05 | 34.02±2.87 | 0.063 |
| **WC(cm)** | 108.45±10.67 | 108.74±11.23 | 107.00±8.31 | 0.166 |
| **SM(kg)** | 32.37±5,83 | 32.55±6.21 | 31.48±3.72 | 0.633 |
| **BF(kg)** | 39.03±9.04 | 39.46±9.44 | 36.92±7.17 | 0.188 |
| **PBF(%)** | 5.58±1.14 | 40.34±6.11 | 39.44±4.09 | 0.063 |
| **SBP(mmHg)** | 128±15 | 128±15 | 127±19 | 0.092 |
| **DBP(mmHg)** | 79±12 | 79±13 | 79±12 | 0.059 |
| **FPG(mmol/L)** | 4.87±0.98 | 4.79±0.95 | 5.27±1.17 | 0.192 |
| **HbA1c(%)** | 5.86±0.65 | 5.81±0.66 | 6.14±0.55 | 0.063 |
| **TG(mmol/L)** | 2.35±1.28 | 2.09±1.02 | 3.67±1.75 | 0.076 |
| **TC(mmol/L)** | 4.87±1.00 | 4.89±1.07 | 4.79±0.51 | 0.388 |
| **LDL-C(mmol/L)** | 2.76±0.79 | 2.82±0.86 | 2.47±0.20 | 0.112 |
| **ALT(U/L)** | 51.27±47.73 | 51.62±51.39 | 51.60±21.93 | 0.069 |
| **GGT(U/L)** | 50.67±48.55 | 46.84±44.68 | 79.80±52.39 | 0.120 |
| **ALP(U/L)** | 67.40±17.27 | 65.68±14.81 | 76.00±27.08 | 0.537 |

**Table S2. Comparison of physical characteristics and metabolic profile of EG group at four time points. Values are presented asMean±SD. BW:Body weight; BMI:Body Mass Index; WC:Waist circumference; SM:Skeletal Muscle; BF:Body fat; PBF:Percentage of Body Fat; SBP:Systolic Blood Pressure; DBP:diastolic blood pressure; FPG:Fasting Plasma Glucose; HbA1c:glycosylated hemoglobin; TG:Triglyceride; TC:Total Cholesterol; HDL-C: High Density Lipoprotein-Cholesterol; LDL-C:Low Density Lipoprotein-Cholesterol; AST: Aspartate Aminotransferase; ALT: Alanine Transaminase; GGT: Glutamyl Transpeptidase; ALP: Alkaline Phosphatase; SCR:Serum Creatinine; UA: Uric acid. Baseline, Mid-piont means 1/2 CR, End-point means 1/4 CR, Post-CR means low CR for 1 month. Values are expressed as the mean±standard deviation(M±SD) of the indicated number of replicates. [a]*P* < 0.01 ,[b]*P* < 0.05 compared with the Baseline.**

| Clinica data / Time points | BW (kg) | BMI ($kg/m^2$) | WC (cm) | SM (kg) | BF (kg) | PBF (%) | SBP (mmHg) | DBP (mmHg) | FPG (mmol/L) | HbA1c(% |
|---|---|---|---|---|---|---|---|---|---|---|
| **Baseline** | 97.53± 15.67 | 34.64± 4.05 | 108.74± 11.23 | 32.55± 6.21 | 39.46± 9.44 | 40.34± 6.11 | 128± 15 | 79± 13 | 4.79± 0.95 | 5.81± 0.66 |
| **Mid-point** | 92.91± 15.40 | 32.98± 3.93 | 101.20± 21.83 | 31.66± 6.13 | 36.40± 8.97[a] | 39.05± 6.13 | 122± 11 | 75± 9 | 4.27± 0.44[b] | 5.90± 0.53 |
| **End-point** | 89.92± 14.98[b] | 31.94± 3.95[a] | 102.64± 12.43[b] | 31.00± 5.97 | 34.36± 9.17 | 38.02± 6.61 | 122± 13 | 72± 10[b] | 4.07± 0.41[a] | 5.56± 0.45 |
| **Post-CR** | 88.89± 15.61[b] | 31.52± 3.85[a,] | 101.08± 11.49[b] | 31.20± 6.07 | 32.79± 9.03[b] | 36.66± 6.34 | 119± 13[b] | 73± 11 | 4.50± 0.46 | 5.22± 0.41[a] |

| Clinica data / Time points | TG (mmol/L) | TC (mmol/L) | HDL-C (mmol/L) | LDL-C (mmol/L) | AST (U/L) | ALT (U/L) | GGT (U/L) | ALP (U/L) | SCR (umol/L) | UA (umol/L) |
|---|---|---|---|---|---|---|---|---|---|---|
| **Baseline** | 2.09± 1.02 | 4.89± 1.07 | 1.14± 0.17 | 2.82± 0.86 | 32.40± 25.30 | 51.32± 51.69 | 44.84± 46.68 | 65.68± 14.81 | 65.40± 16.02 | 411.12± 121.30 |
| **Mid-point** | 1.46± 0.40[a] | 4.64± 1.08 | 1.04± 0.16 | 2.75± 0.91 | 38.64± 34.28 | 50.16± 40.04 | 29.40± 22.06 | 64.20± 13.39 | 69.80± 14.89 | 479.24± 131.50 |
| **End-point** | 1.63± 0.79 | 4.47± 0.85 | 1.08± 0.15 | 2.55± 0.72[c] | 29.68± 17.60[d] | 41.00± 32.39 | 23.20± 15.40[b] | 60.20± 13.57 | 66.76± 13.28 | 429.68± 124.15 |
| **Post-CR** | 1.63± 0.67[d] | 4.66± 0.89 | 1.11± 0.15 | 2.63± 0.75 | 22.52± 6.99[a,d,e] | 29.40± 23.74[c,e] | 30.62± 20.12[e] | 64.12± 13.77[f] | 67.48± 22.15 | 385.56± 118.30[c,f] |

**Table S3 Summary of results of resting-state fMRI with significant differences and corresponding time points in EG. ReHo analysis was corrected by GRF (P < 0.005), see methods.**

| Time points | Brain region | MNI Coordinates | | | T value |
|---|---|---|---|---|---|
| | | X | Y | Z | |
| mid point-baseline | Left orbital inferior frontal gyrus | -15 | 15 | -24 | -4.246 |
| end point-baseline | Putamen | 30 | -21 | 0 | -4.714 |
| post CR-baseline | Right putamen | 27 | 18 | 9 | -4.474 |
| | Left orbital inferior frontal gyrus | 27 | 30 | -9 | -5.112 |
| | Anterior cingulate cortex | 15 | 15 | 36 | -6.425 |
| | Left dorsolateral prefrontal cortex | -12 | 21 | 45 | -5.267 |

**Table S4. Summary of differentially modified modification sites (modified proteins) between CR and obese rats. Proteins were filtered with a threshold value of expression fold change >1.2 and P < 0.05 (Fisher' s exact test).**

| parameter | Type | Up-regulated | Down-regulated |
|---|---|---|---|
| Global proteomes | | 41 | 32 |
| Phospho-proteomes | Sites | 198 | 172 |
| | Proteins | 149 | 145 |
| Acetyl-proteomes | Sites | 32 | 24 |
| | Proteins | 22 | 21 |

**Table S5. Summary of GO function annotations for proteins with significant differences in phosphorylated modification sites (two-tailed Fisher' s exact test to test).**

| GO Terms Level 1 | GO Terms Level 2 | No. of Protein |
|---|---|---|
| Biological Process | cellular process | 188 |
| | biological regulation | 163 |
| | single-organism process | 160 |
| | metabolic process | 90 |
| | multicellular organismal process | 87 |
| | response to stimulus | 86 |
| | cellular component organizatio... | 82 |
| | developmental process | 80 |
| | localization | 73 |
| | signaling | 64 |
| | other | 60 |
| Cellular Component | cell | 206 |
| | organelle | 159 |
| | membrane | 121 |
| | macromolecular complex | 67 |
| | synapse | 47 |
| | cell junction | 38 |
| | membrane-enclosed lumen | 38 |
| | extracellular region | 32 |
| | supramolecular complex | 25 |
| | other | 1 |
| Molecular Function | binding | 212 |
| | catalytic activity | 59 |
| | molecular function regulator | 35 |
| | transporter activity | 27 |
| | structural molecule activity | 22 |
| | molecular transducer activity | 12 |
| | signal transducer activity | 11 |
| | other | 9 |

### Supplementary Figures

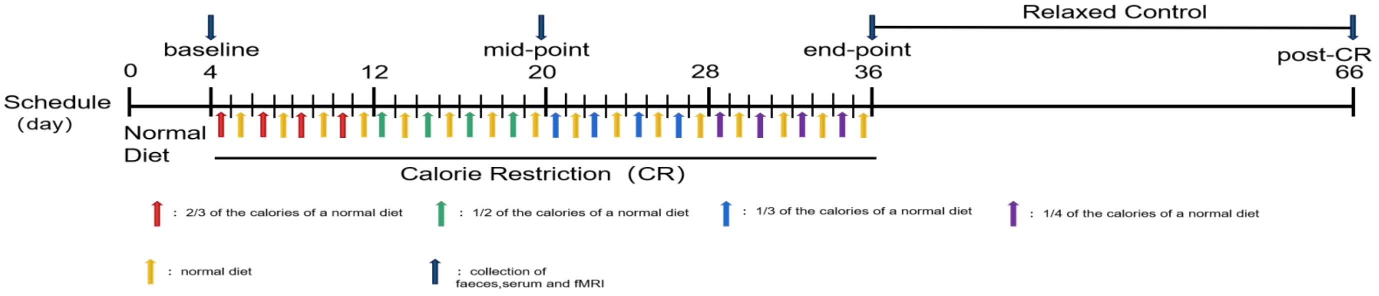


**Figure S1: Schematic experimental outline of the study. The study includes three stages: Normal diet, calorie restriction, and relaxed control. The physiological and biochemical index, feces, serum, and fMRI data were collected at the time of baseline, mid-point, end-point and post-CR shown in the figure.**

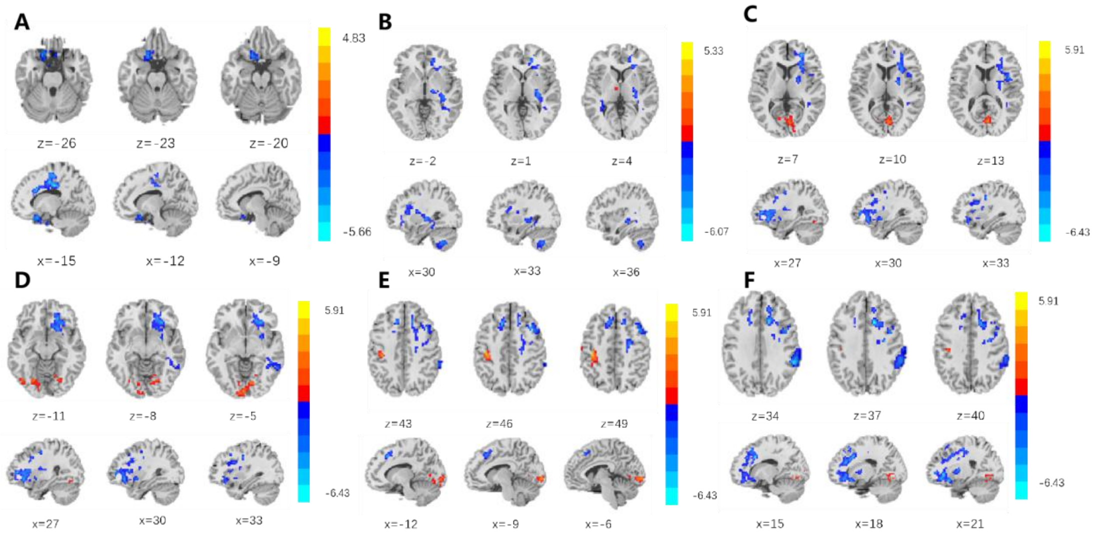


**Figure S2: Graphic representation of results in resting-state fMRI at different time points in EG. ReHo analysis results of rest fMRI in the EG. ReHo analysis was corrected by GRF (P < 0.005), blue color show decreased ReHo value, while red color shows increasd ReHo value). A. mid point *vs*. baseline: Reho value of the OFC decreased. B. end-point *vs*. baselin: Reho value of the putamen decreased. C-F. point-CR *vs*. baseline: Reho value of the OFC, right putamen, left DLPFC, ACC decreased.**

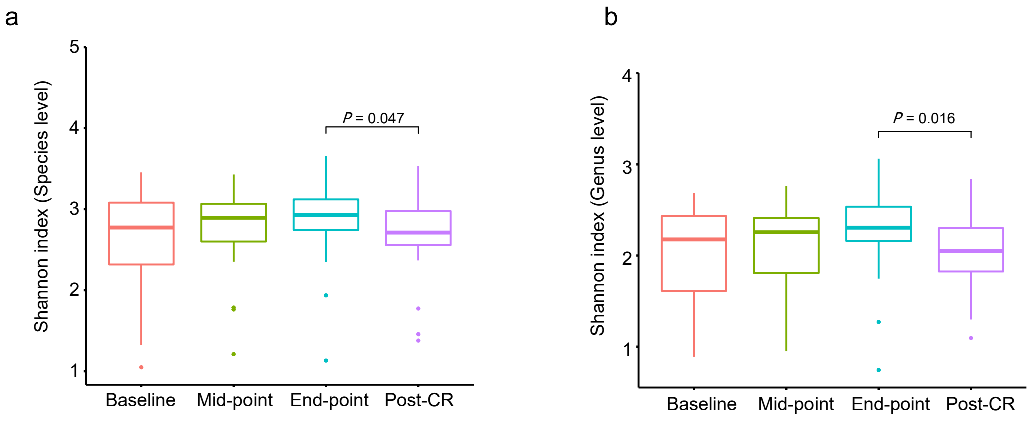


**Figure S3. Alpha diversity in gut microbiome under CR. The Shannon index at baseline, mid-point, end-point and post-CR in species (a) and genus (b) level are shown (Wilcoxon rank-sum test).**

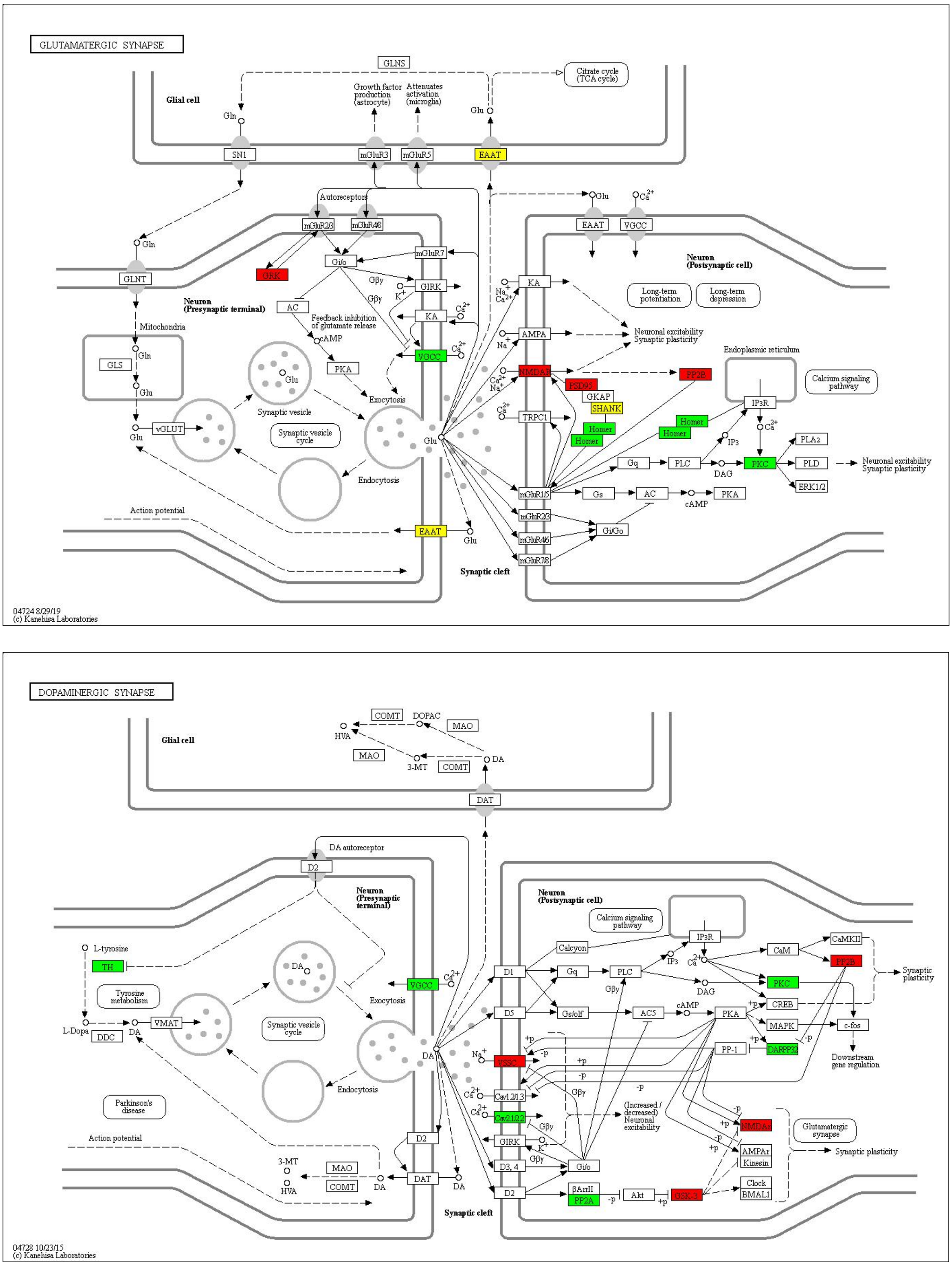


**Figure S4. Enrichment of pathway analysis. Graphical summary of key neural-chemical pathways affected by CR in rats. Glutamatergic synapse and dopaminergic synapse pathway cascades are identified in from the phospho-proteomes data analysis. The proteins in red are upregulated, and the proteins in green are downregulated in CR rats; while yellow indicates that there are multiple proteins in this node and include inconsistently-regulated proteins. a corrected p-value<0.05 was considered significant (two-tailed Fisher's exact test).**
